# Higher performers upregulate brain signal variability in response to more feature-rich visual input

**DOI:** 10.1101/249029

**Authors:** Douglas D. Garrett, Samira Epp, Maike Kleemeyer, Ulman Lindenberger, Thad A. Polk

## Abstract

The extent to which brain responses differ across varying cognitive demands is referred to as “neural differentiation,” and greater neural differentiation has been associated with better cognitive performance in older adults. An emerging approach has examined within-person neural differentiation using moment-to-moment brain signal variability. A number of studies have found that brain signal variability differs by cognitive state; however, the factors that cause signal variability to rise or fall on a given task remain understudied. We hypothesized that top performers would modulate signal variability according to the complexity of sensory input, upregulating variability when processing more feature-rich stimuli. In the current study, 46 older adults passively viewed face stimuli and house stimuli during fMRI. Low-level analyses of our stimuli showed that house images were more feature-rich than faces, and subsequent computational modelling of ventral visual stream responses (HMAX) revealed that houses were more feature-rich especially in V1/V2-like model layers. Notably, we then found that participants exhibiting greater face-to-house upregulation of brain signal variability in V1/V2 (higher for house relative to face stimuli) also exhibited more accurate, faster, and more consistent behavioral performance on a battery of offline visuo-cognitive tasks. Further, control models revealed that face-house modulation of mean brain signal was relatively insensitive to offline cognition, providing further evidence for the importance of brain signal variability for understanding human behavior. We conclude that the ability to align brain signal variability to the complexity of perceptual input may mark heightened trait-level behavioral performance in older adults.

---

The extent to which brain responses differ as a function of cognitive demands is referred to as “neural differentiation,” and has been found to serve as a marker of better cognitive performance (Carp et al. 2010; Park et al. 2010). To date, averaged (mean) brain signals have formed the basis of this line of research. However, an emerging approach has examined within-person neural differentiation using the level of moment-to-moment brain signal variability (i.e., variability-based differentiation, or “VBD”) to characterize different cognitive states. Brain signal variability is generally more sensitive to different cognitive states than are mean brain signals (Garrett, Samanez-Larkin, et al. 2013), and younger and higher performing adults often modulate variability across cognitive states more than do older, poorer performers (Garrett et al. 2010, 2011; Garrett, Kovacevic, et al. 2013; Grady and Garrett 2014; Garrett et al. 2015).

We have hypothesized for some time that the ability to modulate variability across distinct cognitive states should typify organisms able to flexibly and optimally adapt to a host of environmental challenges (Garrett, Samanez-Larkin, *et al*. 2013). Specifically, we and others have postulated that modulation of brain signal variability could reflect differences in stimulus input (Knill and Pouget 2004; Ma et al. 2006; Beck et al. 2008; Garrett, Samanez-Larkin, *et al*. 2013; Orban et al. 2016). For example, early visual regions may be actively suppressed in response to more common stimuli, yet may exhibit a more dynamic response to more differentiated stimuli (Hamm and Yuste 2016; Homann et al. 2017; Vinken et al. 2017). However, evidence for this effect is lacking in humans, as are the associated behavioral consequences. One perspective is that the brain could conceivably limit resource allocation (narrowing neural dynamic range) when stimulus input is more reducible, and it should also upregulate dynamic range to the extent that more resource intensive processing is required (e.g., to encode more differentiated sources of sensory input) (Mlynarski and Hermundstad 2018). Faces and houses, two of the most studied categories of visual stimuli in cognitive neuroscience, provide potentially fertile experimental materials in this context. Humans are considered “face processing experts” (Carey and Diamond 1977; Carey 1992), and faces can be reduced to a limited number of statistical/perceptual dimensions, yet still be processed, discriminated, and recognized (Sirovich and Kirby 1987; O’toole et al. 1993; Tsao and Livingstone 2008). In line with Mlynarski et al. (2018), face processing may in turn require lower encoding/processing fidelity and minimal neural dynamic range because processing demands are lower. Houses, in contrast, are a much more differentiated stimulus type, with relatively few constraints on their form relative to faces, potentially increasing required encoding/processing demands and neural dynamic range (See Figure 1 for a depiction). At its core, VBD (a presumed marker of neural flexibility) may reflect one’s ability to tune to the dynamics of the external world. We expect that those better able to upregulate brain signal variability when processing more complex visual stimulus categories should also exhibit better trait-level cognitive performance, as such individuals would be more adaptable to effectively process different types (or degrees) of stimulus input.

**Figure 1:**
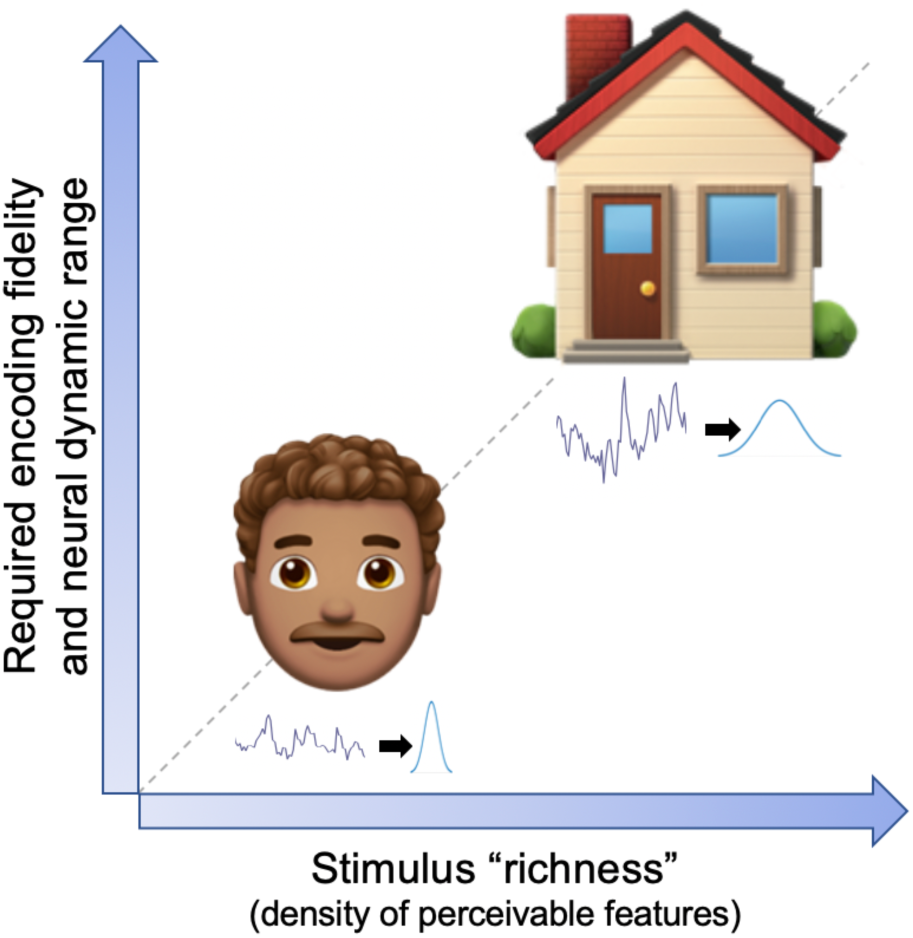
Theoretically, greater stimulus “richness” should require higher encoding fidelity, potentially expressed as higher neural dynamic range (variability). Under the expectation that houses are a more differentiated stimulus category than faces, encoding houses should also require higher neural dynamic range (variability).

In the present study, we tested these various predictions in a dataset in which face and house stimuli were passively viewed by 46 older adults (59-73 years) during fMRI scanning. Under expectations that greater “visual feature content” may upregulate brain signal variability in higher performers, we first quantified differences in visual feature content across face and house image categories using a series of standard stimulus image processing metrics (e.g., pixel-based standard deviation, entropy). However, because such image differences *per se* don’t provide biologically-informed predictions of how those differences may manifest in the brain, we also utilized a computational model of the ventral visual stream (HMAX) to help isolate the visuo-cortical regions potentially most sensitive to face/house content differences. HMAX estimates the presence of simple and composite visual features in the stimuli it is fed; if the model detects more visual features for a given stimulus category, then we consider that category more “feature-rich.” In this way, we leverage HMAX as a simple, yet neurally-inspired estimator of feature-richness in the exact stimuli seen by our participants in scanner. Crucially, using multivariate modeling, we then tested whether these visual region-specific HMAX effects may converge with those brain regions in which greater VBD between faces and houses during fMRI was associated with better trait-level cognitive performance. Finally, we compared VBD to standard mean signal-based face-house modulation to gauge relative sensitivity of each brain measure to latent-level behavioral outcomes.

## MATERIALS and METHODS

### fMRI experiment

#### Participants

The initial sample utilized in the current study consisted of 48 community-dwelling older adults aged 59-73 (mean 66.05 ± 4.40, 30 female), and represent the pre-training sample from a previously described study (Kleemeyer et al. 2016; Kleemeyer et al. 2017). All participants had MMSE score ≥ 26, were free of neurological, psychiatric, and cardiovascular diseases, were right-handed, and were suitable for MR assessment (e.g. no magnetic implants, no claustrophobia). This study was carried out in accordance with the recommendations of the ethics committee of the German Psychological Society. All participants gave written informed consent in accordance with the Declaration of Helsinki and participated voluntarily. They were paid for study completion. Two subjects were discarded from the current sample, one with improper slice positioning during scanning, and another was an extreme outlier on the Stroop task (an offline speeded measure; see below). Thus, analyses were based primarily on data from 46 participants.

#### MRI Data Acquisition, Task, and Preprocessing

Brain images were acquired on a Siemens TIM Trio 3T MRI scanner (Siemens, Erlangen, Germany) at the Max Planck Institute for Human Development in Berlin. A high-resolution T1-weighted MPRAGE (TR = 2500 ms, TE = 4.76 ms, TI = 1100 ms, flip angle = 7°, acquisition matrix = 256 × 256 × 176, 1 mm isotropic voxels) was first acquired. A conventional echo-planar MR sequence was then used for functional acquisitions (TR = 2000 ms, TE = 30 ms, flip angle = 80°, FOV = 216mm) encompassing 192 volumes per run and 36 slices per volume (slice thickness 3 mm).

During functional imaging, participants passively viewed greyscale face, house, or phase-scrambled face/house images (following the procedures of Park *et al*. (2010)) across two runs, each of which consisted of four blocks per stimulus category. The face pictures were taken from previous work (Minear and Park 2004). The house pictures were digital photographs of houses in the Ann Arbor, Michigan area. Each block contained 15 images shown for 2 seconds each, resulting in 30 second block lengths (total of 6 minutes per run). Stimuli were presented via E-prime (Psychology Software Tools, Pittsburgh, PA) and displayed by a projection system.

fMRI data were preprocessed with FSL 5 (RRID:SCR_002823) (Smith et al. 2004; Jenkinson et al. 2012). Pre-processing included motion-correction with spatial smoothing (7 mm full-width at half maximum Gaussian kernel) and bandpass filtering (.01-.10 Hz). We registered functional images to participant-specific T1 images, and from T1 to 2mm standard space (MNI 152_T1) using FLIRT. We then masked the functional data with the GM tissue prior provided in FSL (thresholded at probability > 0.37). We detrended the data (up to a quadratic trend) using the SPM_detrend function in SPM8. We also utilized extended preprocessing steps to further reduce data artifacts (Garrett *et al*. 2010, 2011; Garrett *et al*. 2015). Specifically, we examined all functional volumes for artifacts via independent component analysis (ICA) within-run, within-person, as implemented in FSL/MELODIC (Beckmann and Smith 2004). Noise components were identified according to several key criteria: a) Spiking (components dominated by abrupt time series spikes); b) Motion (prominent edge or “ringing” effects, sometimes [but not always] accompanied by large time series spikes); c) Susceptibility and flow artifacts (prominent air-tissue boundary or sinus activation; typically represents cardio/respiratory effects); d) White matter (WM) and ventricle activation {Birn:2012gh}; e) Low-frequency signal drift (Smith et al. 1999); f) High power in high-frequency ranges unlikely to represent neural activity (≥ 75% of total spectral power present above .10 Hz;); and g) Spatial distribution (“spotty” or “speckled” spatial pattern that appears scattered randomly across ≥ 25% of the brain, with few if any clusters with ≥ 80 contiguous voxels [at 2×2×2 mm voxel size]). Examples of various components we typically deem to be noise can be found in Garrett et al. (2014). By default, we utilized a conservative set of rejection criteria; if manual classification decisions were challenging due to mixing of “signal” and “noise” in a single component, we generally elected to keep such components. Three independent raters of noise components were utilized; > 90% inter-rater reliability was required on separate data before denoising decisions were made on the current data. To enable semi-automated data denoising using FSL FIX, we manually classified 30% of participant data to provide a noise component training set. Features from the noise component training set were then extracted and used to detect noise components from the remaining 70% of participant data via FIX. Upon evaluating the automated labelling for several subjects against our manual decisions, we used a FIX threshold of 60, which permitted a best match to manual decisions of two independent raters. Components identified as artifacts were then regressed from corresponding fMRI runs using the *regfilt* command in FSL. We found that these additional preprocessing steps had dramatic effects on the predictive power of SD_BOLD_ in past research, effectively removing 50% of the variance still present after traditional preprocessing steps, while simultaneously doubling the predictive power of SD_BOLD_ (Garrett *et al*. 2010). Critically, our recent work also suggests that when such denoising approaches are applied, age differences in SD_BOLD_ remain robust to multiple vascular controls measured via dual-echo ASL-BOLD using carbogen-based hypercapnia (Garrett et al. 2017). The present sample only contains a narrow age range of older adults (59-73 years), further minimizing the potential impact of aging-based differences in vasculature. For all robust models presented in the current paper, we also partial chronological age from all effects to further confirm that any residual (post-denoising) age-related artifacts are controlled (see Results).

#### Voxel-wise estimates of SD_BOLD_ and comparison to mean_BOLD_

To calculate SD_BOLD_, we first performed a block normalization procedure to account for residual low frequency artifacts. We normalized all blocks for each condition such that the overall 4D mean across brain and block was 100. For each voxel, we then subtracted the block mean and concatenated across all blocks. Finally, we calculated voxel standard deviations across this concatenated time series (Garrett *et al*. 2010). All models described below were run on grey matter (GM) only, after a standard GM mask derived from the MNI152 average brain was applied to each 4D image set.

We sought also to compare SD_BOLD_ results to a typical mean-based measure of BOLD activity (mean_BOLD_). We calculated mean signal (mean_BOLD_) for each experimental condition as follows; we first expressed each signal value as a percent change from the average of the last four scans from the previous block, and then calculated a mean percent change within each block and averaged across all blocks for a given condition (a typical method in the PLS data-analysis framework). This effectively acts as an explicit high-pass filter over the data. We then re-ran relevant PLS models described below, while using mean_BOLD_ measures.

### Offline visuo-cognitive assessment

Participants completed a 90-minute cognitive testing session outside the scanner (see Kleemeyer, 2018). For the current study, the available cognitive battery spanned working memory/updating, reasoning, inhibition, and switching. Responses for computerized tasks were provided via response boxes or a computer keyboard. For the current study, eight tasks were included, four tasks in which accuracy was emphasized and four in which speed was emphasized. For all tasks, if initial inspection of normality of the between-subject distribution failed, we attempted to transform to Gaussian via either log or square root transformation (and confirmed success with a combination of Kolmogorov-Smirnov tests for normality and Q-Q plots).

#### Accuracy-based measures

We assessed working memory/updating using three different tasks: letter updating, spatial updating and 2-back (all taken from Schmiedek et al. (2009)). During letter updating, single letters (A, B, C, or D) appeared randomly on the screen one after the other for 2500 ms with a 500 ms ISI. The presentation sequence stopped after 7, 9, 11, or 13 letters and the task instructed participants to enter the last three letters displayed, via the computer keyboard. After four practice trials, participants completed two sequences composed of 7, 9, 11, or 13 letters each, resulting in a total of eight trials presented in randomized order. During spatial updating, participants were shown two 3 × 3 grids, presented side-by-side on the computer screen. At the beginning of every trial, a dot appeared simultaneously in each grid for 4000 ms and participants were instructed to remember the position of the dot. Next, arrows appeared synchronously above each grid for 2500 ms, indicating that the dot in the respective grid had to be moved one field in the direction of the arrow. After 500 ms another pair of arrows required another moving of the dots. At the end, the final position of the dots had to be marked in the grids via mouse click. After a practice trial, participants completed two easy trials including two updating operations per grid, as well as two difficult trials including three updating operations per grid. During the 2-back, single digits (1-9) appeared randomly on the screen for 500 ms one after the other with a ISI of 3000 ms. Participants were asked to indicate, whether the currently presented digit was identical to the digit presented two displays earlier in the sequence or not. After one practice sequence (26 numbers), participants completed three test sequences (39 numbers).

Finally, reasoning was assessed using a version of Raven’s progressive matrices (Raven et al. 1998). Participants saw a 3 × 3 matrix with patterns following certain regularities. The pattern on the lower right was missing, and participants were instructed to identify the correct pattern out of eight given alternatives. A total of 15 trials could be completed within a maximum of 15 minutes.

For all four tasks, accuracy served as the outcome measure.

#### RT-based measures

For each measure, all trials in which RT was < 200 ms or greater than ±3 SDs from within-person means were dropped prior to computing reaction time means (RT_mean_) and SDs (RT_sd_) for each subject. RT measures were only computed for correct trials for all tasks. We employed four speeded tasks in total (see below).

Inhibition was assessed via the Stroop task (Stroop, 1935); color-words were shown in either the same (congruent) or different (incongruent) colored font in the center of the screen for 1000 ms. Participants were asked to indicate as quickly as possible the font color; the next stimulus only appeared 1000 ms after an answer was given. Participants completed 24 practice trials and four blocks of 36 test trials each; 50% of trials were incongruent (color-word and font color did not match). Reaction time values for this task were considered for congruent and incongruent trials separately.

Task switching was assessed using three tasks: the number-letter task (Schmitter-Edgecombe and Langill 2006), the global-local task (Kinchla et al. 1983) and the face-word task (Yeung et al. 2006). During the number-letter task, participants saw a number-letter pair appearing in one of four quadrants of the screen (top left, top right, bottom left, bottom right). In cases where the stimulus pair appeared at the top, participants had to attend to the number, and indicate whether it was odd or even. If the stimulus pair appeared at the bottom, participants were instructed to attend to the letter, and indicate whether it was a vowel or a consonant. Stimuli included 2, 3, 4, 5, 6, 7, 8, 9, A, E, I, U, G, K, M, R. For the global-local task, participants were presented with Navon figures, i.e. large objects composed of small objects. Objects were always circles, triangles, squares, or crosses, e.g. a large circle composed of small triangles. If the objects appeared in blue, participants had to indicate the shape of the larger (global) object. If the objects appeared in black, participants had to indicate the shape of the smaller (local) objects. During the face-word task, participants saw 1-or 2-syllable words overlaid on male or female faces. A key appeared below the stimulus pair, indicating whether the face or the word should be attended. For words, participants were required to indicate whether the word had one or two syllables, whereas for faces they had to decide whether it was female or male. For all task-switching tasks, stimuli were presented for 2500 ms and the next stimulus only appeared 500 ms after an answer was given via response boxes. Trials were randomly presented, with 50 % of the trials requiring a task-switch. For every task, participants completed 24 practice trials followed by 128 test trials, divided into four separate blocks of 32 trials each. Reaction time values for these tasks were considered for switch and non-switch trials separately.

For each of the four tasks, RT_mean_ and RT_SD_ served as metrics of interest.

### Statistical modeling: Partial Least Squares

To examine a multivariate contrast of SD_BOLD_ across face and house conditions, we employed a Task partial least squares (PLS) analysis (Mcintosh et al. 1996; Krishnan et al. 2011). Task PLS begins by calculating a between-subject covariance matrix (COV) between experimental conditions/groups and each voxel’s SD_BOLD_, which is then decomposed using singular value decomposition (SVD). This yields a left singular vector of experimental condition/group weights (*U*), a right singular vector of brain voxel weights (*V*), and a diagonal matrix of singular values (*S*). Task PLS produces orthogonal latent variables (LVs) that optimally represent relations between experimental conditions/groups and voxel-wise SD_BOLD_ values.

To examine multivariate relations between SD_BOLD_ face-house upregulation and overall offline cognitive performance, we utilized behavioral PLS analysis (Mcintosh *et al*. 1996; Krishnan *et al*. 2011). This analysis begins by calculating a between-subject correlation matrix (CORR) between (1) each voxel’s SD_BOLD_ upregulation value (i.e., house SD_BOLD_ minus face SD_BOLD_) and (2) a series of offline cognitive measures (all RT and accuracy-based measures noted above). CORR is then decomposed using singular value decomposition (SVD).

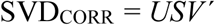

This decomposition produces a matrix of left singular vectors of cognition weights (*U*), a matrix of right singular vectors of brain voxel weights (*V*), and a diagonal matrix of singular values (*S*). For each LV (ordered strongest to weakest in *S*), what results is a spatial activity pattern depicting the brain regions that show the strongest available relation to offline performance. Significance of detected relations of both PLS model types was assessed using 10000 permutation tests of the singular value corresponding to the LV. A subsequent bootstrapping procedure revealed the robustness of within-LV voxel saliences across 10000 bootstrapped resamples of the data (Efron and Tibshirani 1993). By dividing each voxel’s weight (from *V*) by its bootstrapped standard error, we obtained “bootstrap ratios” (BSRs) as normalized estimates of robustness. For the whole brain analysis, we thresholded BSRs at values of ±3.00 (∼99.9% confidence interval).

We also obtained a summary measure of each participant’s robust expression of a particular LV’s spatial pattern (a within-person “brain score”) by either (1) multiplying the vector of brain weights (*V)* from each LV by within-subject vectors of voxel SD_BOLD_ values (separately for each condition/group within person) for the Task PLS models, or (2) in the behavioral PLS model noted above, multiplying the model-based vector of voxel weights (*V*) by each subject’s vector of voxel SD_BOLD_ upregulation values (*P*), producing a single within-subject value:

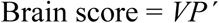

Similarly, for our behavioral PLS model linking brain and behavior, we also computed a latent “behavior score” for each subject by multiplying the model-based vector of behavior weights (*U*) by each subject’s vector of values on the behavioral tasks (*Q*):

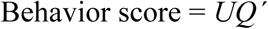

By then correlating Brain and Behavior scores, latent level relations can be plotted (see Results).

### Analysis of face/house stimulus properties

#### Image metrics

In the current fMRI task design, participants passively viewed face and house stimuli in several blocks of trials. To better characterize lower level stimulus differences between the current face and house stimuli, we computed various image metrics estimable via the Image Processing Toolbox in Matlab 2018a: (1) <u>contrast (or “variance”)</u>: the intensity contrast between neighbouring pixels over an entire 2D image (contrast = 0 for a constant image); (2) <u>correlation</u>: correlation between neighbouring pixel intensities over a 2D image; (3) <u>energy (or “inertia”)</u>: the sum of squared elements in the grey-level co-occurrence matrix (GLCM; energy = 1 for a constant image; (4) <u>homogeneity</u>: measure of the closeness of the distribution of elements in the GLCM to the GLCM diagonal (homogeneity = 1 for a constant image); (5) <u>local entropy (using</u> *<u>entropyfilt</u>*): This function computes entropy on greyscale intensities from 9×9 pixel neighbourhoods, within-image; we then averaged all local neighbourhoods within-image to estimate a single average image entropy per face and house image. Estimating and averaging local image intensity entropy in this manner allows the algorithm to detect mean-level predictability of surrounding intensities from any given intensity within the image; (6) <u>local standard deviation (</u>*<u>Stdfilt</u>*): using the same grey scale intensity neighbourhood approach as above (9×9 pixel), this function computes the standard deviation (SD) for each neighborhood; we then averaged all local SD neighbourhoods within-image to estimate a single average image SD per face and house image. All individuals received the same face and house stimuli, so all metrics for image properties shown in the current paper were the same for all participants by definition.

#### Estimation of visuo-cortical layer-specific stimulus feature differentiation via a feedforward computational model of the ventral visual system (HMAX)

Given that all participants saw the same face and house images, we elected to present each image to the HMAX feedforward model of the ventral visual system to quantify which visuo-cortical model layers may be most sensitive to differences between the exact face and house images seen by our participants (Riesenhuber and Poggio 1999; Serre, Oliva, et al. 2007; Serre, Wolf, et al. 2007) (code is freely available at: http://maxlab.neuro.georgetown.edu/hmax.html). HMAX consists of four layers (see Fig 2). Layers S1 and C1 correspond to V1/V2 function, and layers S2 and C2 to V2/V4 function (Serre et al. 2005). Within the first layer (S1), a range of Gabor filters (intended to provide a principled model of cortical simple cell receptive fields, with 16 different filters corresponding to *n* x *n* pixel neighborhoods (sizes: [7:2:37])) and four orientations (HMAX defaults: −45°, 0°, 45°, 90°) are fitted to each image in overlapping windows (50% overlap). The resulting S1 map corresponds to simple cell responses for all positions within the input image, and the fitting procedure is completed for each orientation and filter size separately.

**Figure 2:**
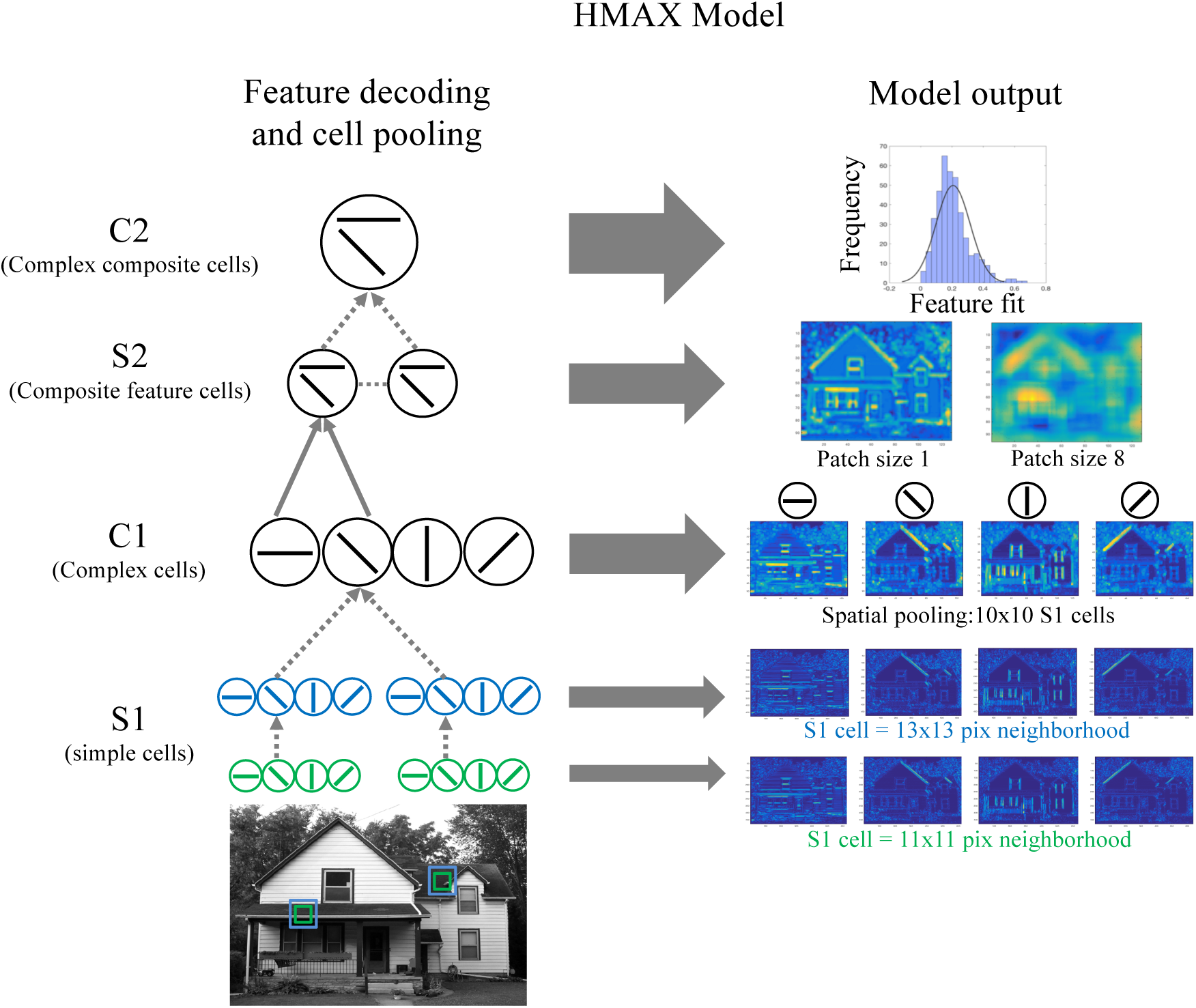
Visualization of the HMAX model of ventral visual cortex (see Riesenhuber et al. (1999) and Serre et al. (2007)). Left panel (“Feature decoding and cell pooling”): the model is sketched. Two example house image locations and neighborhood sizes are shown (green/blue boxes), from which all orientations are decoded. At the S1 (simple cell) layer, these locations are convolved with different Gabor filter sizes within each orientation (here, two filter (neighborhood) sizes of 11×11 (in green) and 13×13 pixels (in blue), and 4 orientations, are shown). At the C1 layer, the model takes the max (dotted paths) simultaneously over both filter sizes and over a pool of S1 cells (here, 10×10 cells) for each orientation separately. The S2 layer represents hybrid features estimated via weighted sum (solid grey paths) after comparison of image features to 400 prototypical features from an external image set, separately for a series of C1 cell neighborhood sizes within each scale band (example scale band 2 is shown). Finally, layer C2 represents the max taken over all S2 cells to generate an aggregated “fit” (inverse Euclidean distance) of image features to the 400 feature prototypes. Right panel (“Model output”): The output of these example steps for the entire image is shown, with warmer colors representing better fits to each orientation (layers S1 and C1) and prototype (layer S2).

At the next layer (complex cells in C1), two consecutive maximization steps over S1 simple cells are computed: (a) over two neighboring filter sizes for each S1 cell, and then (b) over a pool of S1 cells. Because 16 filter sizes are used in HMAX [7:2:37 pixels], taking the max over neighboring pairs of filters results in eight “scale bands”. The scale band index then determines the spatial neighborhood of S1 cells over which outputs are pooled in the second step [8:2:22 S1 cells], independently for each orientation (Serre, Wolf, *et al*. 2007). We took the median C1 value within each image, for each scale band and orientation separately. We then compared these within-image median values, within each scale band and orientation, between face and house stimuli. This resulted in 8×4 independent sample t-tests (equal variances not assumed; see Results).

At the third layer (S2, composite feature cells), a template-based approach is used to estimate the fit to simple and complex (hybrid) C1-level prototypical feature sets included in the HMAX code, and derived from a library of naturalistic stimuli (Serre, Wolf, *et al*. 2007). Eight different C1 neighborhood sizes [2:2:16] for each of 400 different features were fitted. S2 cells pool over C1 cells within each neighborhood and scale band (but across all orientations), quantifying the fit between the spatial features of the input image and those of the stored prototypes. The final layer (C2, complex composite feature cells) then takes the max over all scale bands, quantifying the global fit (inverse Euclidean distance) between the features of each image and each prototype, separately for each prototype C1 neighborhood size. When comparing face and house stimuli, we took the median fit per image for each C1 neighborhood size separately, resulting in eight independent samples t-tests (equal variances not assumed; see Results).

Openly available code for all analyses steps will be available on https://github.com/LNDG.

## RESULTS

### Quantifying visual features in face/house images via pixel-based metrics

The current study utilizes face and house images from previously published work, but with a new older-adult-only fMRI sample. However, face and house stimuli may differ along a variety of dimensions. Houses are intuitively more differentiated than faces, with far fewer constraints on their form (e.g., faces have two eyes, but a house could have any number of windows). To examine simple image statistics in the current set of face and house stimuli, we computed several 2-D image metrics: (1) contrast; (2) correlation; (3) energy; (4) homogeneity; (5) entropy (using *entropyfilt*), and (6) standard deviation (using stdfilt; see Fig 3 and Methods for details). It is clear in Fig 3 that houses differ appreciably from faces in nearly every category, and in particular, are much more differentiated images (e.g., higher entropy, lower homogeneity, higher contrast variance).

**Figure 3:**
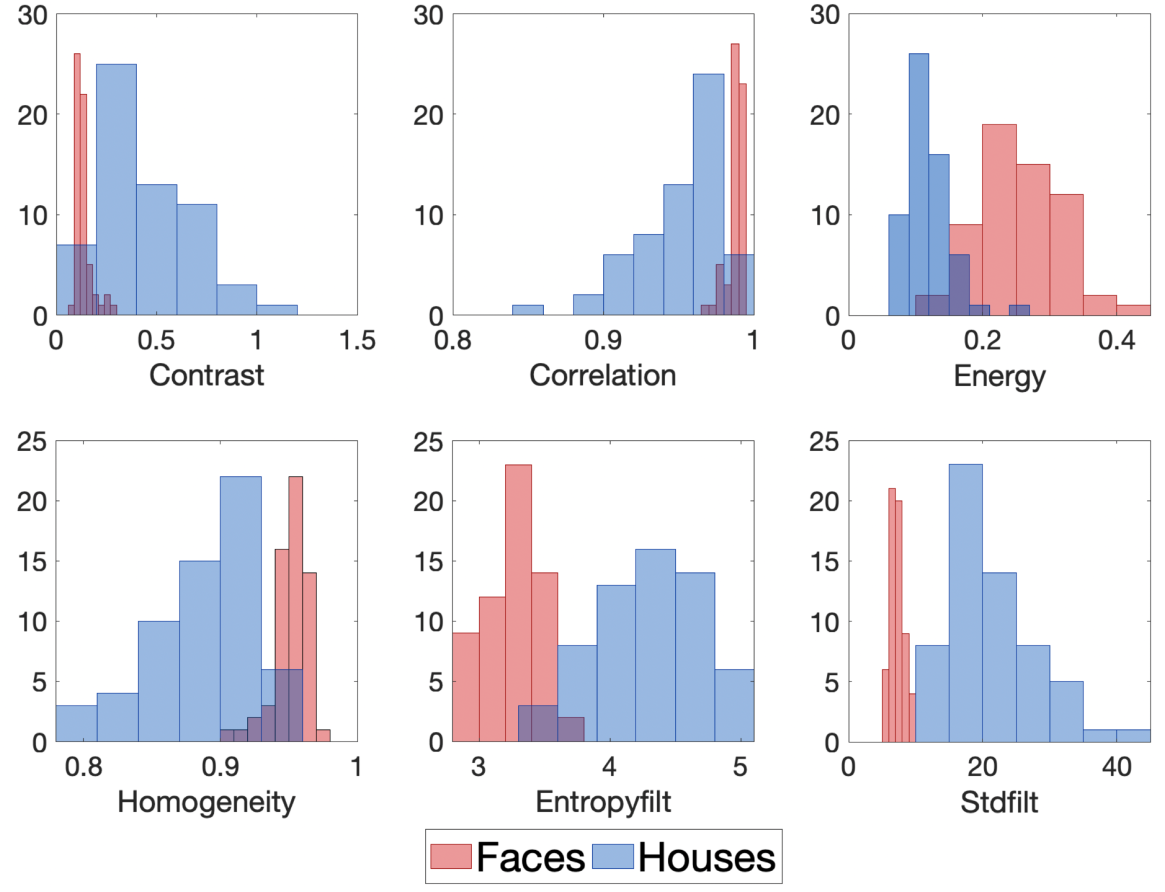
Image metric histograms for face (blue bars) and house (red bars) stimuli. 60 unique images per condition were utilized.

### Quantifying visual features via feedforward computational modeling of the ventral visual stream (HMAX)

Although useful, such pixel-based image category differences *per se* don’t aid biological understanding of how those differences may manifest in the brain. Would our particular face and house images be maximally differentially expressed in V1 or V4, for example? To thus estimate “features” present within our face/house stimuli in potentially sensitive nodes of the ventral visual system, we used a biologically-inspired, openly available feedforward computational model (HMAX) (Riesenhuber and Poggio 1999; Serre *et al*. 2005; Serre, Oliva, *et al*. 2007; Serre, Wolf, *et al*. 2007). See Fig 2 and Methods for details. In short, HMAX estimates the presence of simple and composite visual features in the stimuli it is fed; if HMAX detects more visual features for a given stimulus category, then we consider that category more “feature-rich.” In this way, we are using HMAX as a simple, yet neurally-inspired, estimator of feature differentiation in the stimuli seen by our participants in scanner. Further, because HMAX is not based on the BOLD signal *per se*, it does not deterministically make BOLD-based predictions, nor does it know how BOLD may reflect individual differences in behavior (we will test this in Results below); HMAX will only provide an estimate of visuo-cortical regions sensitive to feature differences in our exact stimuli sets (faces and houses).

We focused our analyses on C1 and C2 layers as aggregate representations of “simple” (V1/V2) and “composite” feature cells (V2/V4) (Serre *et al*. 2005). HMAX model results indicated that each spatial orientation occurred significantly more often for house than for face stimuli across all “receptive field sizes” (i.e. Scales; see the left panels of Figures 4 (for example orientation) and S1 (for all orientations)). Interestingly, condition separation at C1 appeared to increase with increasing scale band (receptive field) size, suggesting that face features may be more “reducible” relative to houses as receptive field size grows in V1/V2 cells (Figure 5). As at layer C1, we similarly found at C2 that house stimuli showed significantly higher median fits (i.e., lower Euclidian distance) to prototypical spatial features across different neighborhood sizes (Figures 4, S1, and 5, right panels). Overall, these findings indicate that our house stimuli are much more feature-rich than our face stimuli. Interestingly, although all *t*-tests exhibited strong effects, the degree of statistical differentiation between face and house conditions was greater for layer C1 (V1/V2-like) than for C2 (V2/V4-like). This highlights that primary/secondary visual cortex (V1/V2) may be particularly sensitive to such condition differences in visual feature density, and serve as target regions in our brain-behavior models below.

**Figure 4:**
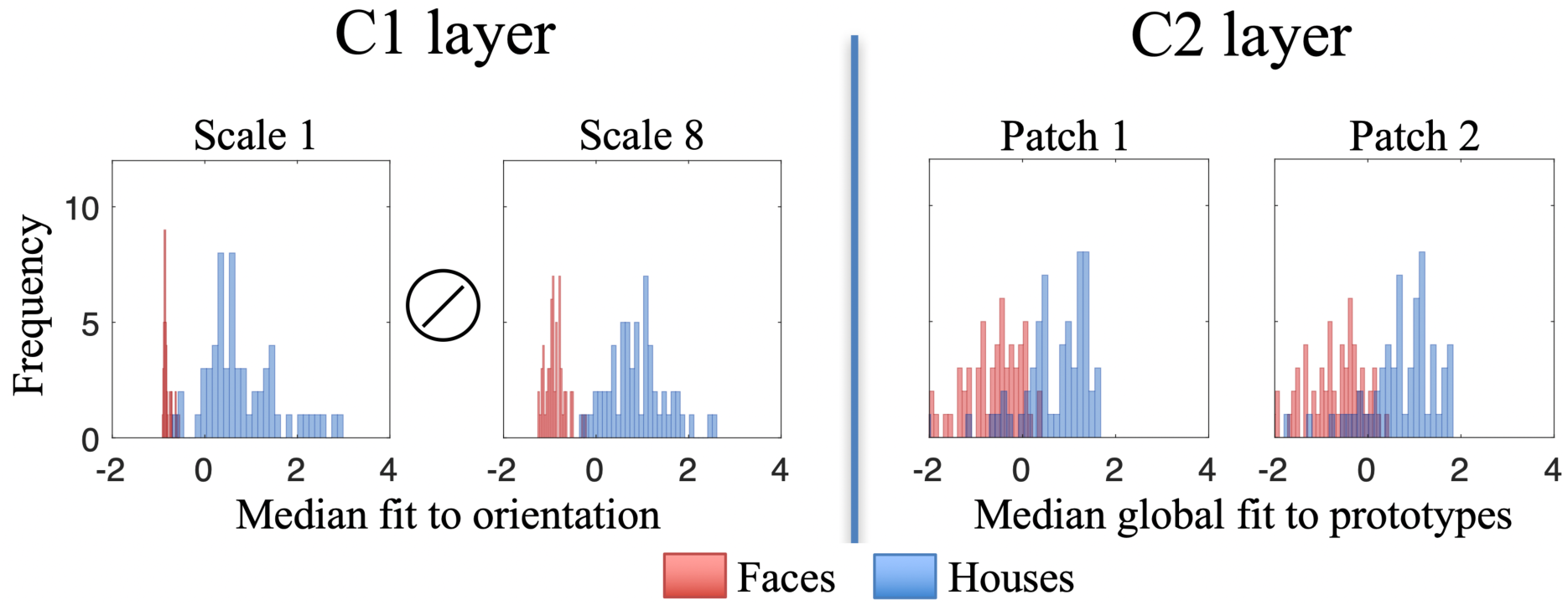
Example HMAX C1 (simple feature) and C2 (composite feature) distributions for face and house stimuli. All 60 unique images per condition from the fMRI stimulus set were submitted to the model. At the C1 layer (left), the median of each C1 map is shown, averaging over all locations within-image, for the smallest and largest receptive field sizes (i.e., scale bands 1 and 8) for an example spatial orientation. At the C2 layer (right), the median within-image fit to a library of 400 feature prototypes is displayed for two exemplar receptive field (or “patch”) sizes. Higher values for C1 and C2 indicate higher “feature richness.” All C1 and C2 value ranges (xaxes) are z-normalized.

**Figure 5:**
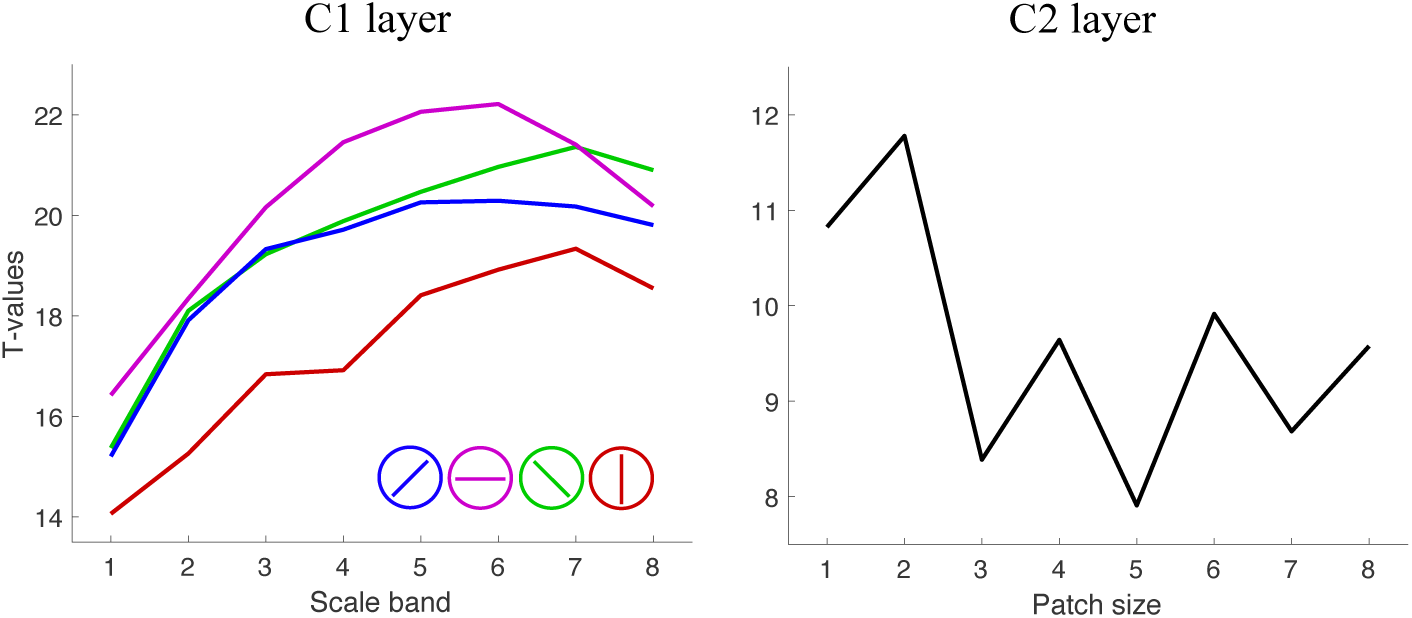
Independent sample t-test values for C1 (V1/V2) and C2 (V2/V4) layer feature richness. All t-values represent house minus face conditions. For C1, *t*-values are plotted *for each orientation and scale band* (associated *p*-value range = ∼1.00* 10^−22^ to 10^−32^). For C2, values are plotted for each patch size (associated *p*-value range = ∼1.00* 10^−12^ to 10^−21^). Equal variances not assumed for any test.

### Multivariate models linking VBD to offline behavior

Next, we examined whether greater within-person upregulation of brain signal variability was linked to better visuo-cognitive performance outside the scanner. In so doing, we attempt to establish trait-based, latent-level relations between behavioral performance and one’s ability to upregulate brain signal variability, in line with presumably more processing intensive and feature-rich sensory input (i.e., for houses more than faces). Notably, although general condition differences in SD_BOLD_ are in line with expectations (i.e., houses evoked more variability than faces overall; see Supp Results and Fig S2), we directly tested the link between face-house differences in variability and offline performance given that the regions expressing maximum differences between faces and houses are not necessarily those that are the most sensitive to individual differences in behavior. To test this, we ran a multivariate partial least squares (PLS) model linking upregulation of SD_BOLD_ from face to house to cognition on a series of 20 measures (four accuracy, eight RT_mean_, eight RT_SD_) across 8 tasks outside the scanner. A robust latent variable (permuted *p* = 0.05, accounting for 59.45% of the crossblock covariance) resulted, indicating that greater face to house upregulation of SD_BOLD_ correlated with more accurate, faster, and more stable offline visuo-spatial performance (latent *r* = 0.47; see Figure 6). This effect was bootstrap robust for 3/4 (accuracy), 8/8 (RT_mean_), and 7/8 (RT_SD_) cognitive measures (see Figure S3) contributing to the overall latent effect.

**Figure 6:**
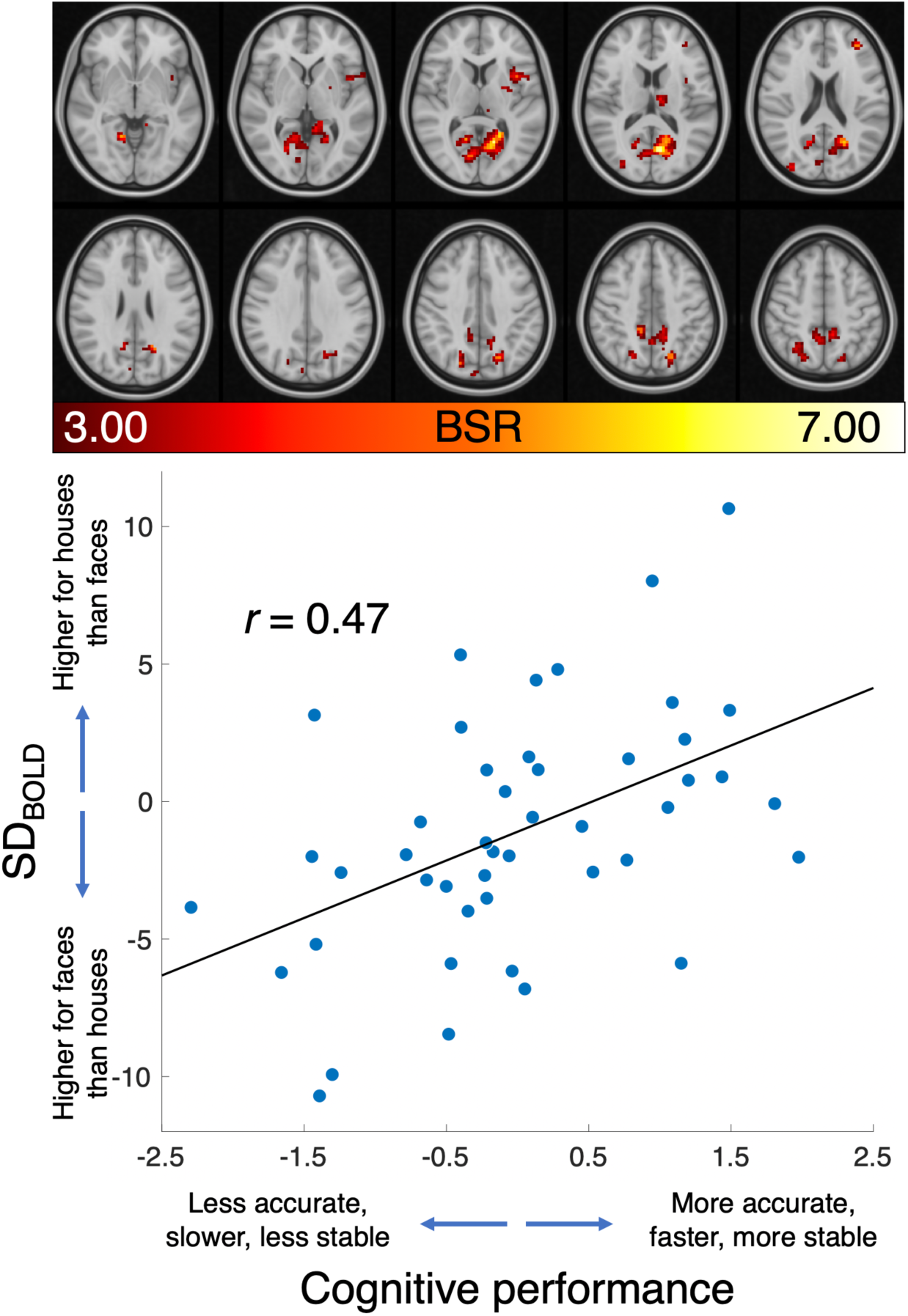
Multivariate model showing greater SD_BOLD_ upregulation during house compared to face conditions (house_SD_BOLD_ minus face_SD_BOLD_) as a function of more accurate, faster, and more stable offline cognitive performance. BSR = bootstrap ratio. Slices increase from Z=-4 (top left) in 6 mm increments.

Spatially, in line with HMAX results noted above, this multivariate effect was most prominent in primary (V1/V2) visual cortex (see Figure 6 and Table S1). Precuneus/posterior cingulate regions were also relevant, as (to a lesser extent) was superior occipital cortex. Interestingly, our recent work has shown that thalamus is particularly central to how the brain fluctuates and communicates at rest (Garrett et al. 2018); here we show first evidence that modulation of thalamic variability also reflects better latent-level cognition. Finally, given that adult age remains a major correlate of individual differences in SD_BOLD_ in previous studies (Garrett, Samanez-Larkin, *et al*. 2013; Grady and Garrett 2014), we also tested the above link between face-house upregulation of SD_BOLD_ and cognitive performance, while controlling for chronological age (59-73 years); the latent level relation noted in Figure 6 between cognitive performance and SD_BOLD_ modulation remained identical (*r*semi-partial = .47), likely due to the use of an older-adult-only sample in the current study.

### Mean_BOLD_-based face-house modulation is insensitive to offline performance

Our previous work shows that SD_BOLD_ is typically more sensitive to individual differences in cognition than is mean_BOLD_ (Garrett *et al*. 2010, 2011; Garrett *et al*. 2014; Garrett *et al*. 2015). To test whether this was also the case in the present study, we used the same types of PLS models as above, instead using voxel-wise mean_BOLD_ as the brain measure of interest. First, a task PLS model of the face-house condition differences (Fig S4) was significant (permuted *p* <.0001), showing that the face condition activated classic regions such as bilateral fusiform face area (FFA), and the house condition activated expected regions such as bilateral parahippocampal place area (PPA), bilateral lingual gyrus, and lateral occipital complex (LOC) (for example and corresponding regional coordinates, see Table 2 in (Mckeeff and Tong 2007)). However, of primary interest was a behavioral PLS model linking face-house condition differences in mean_BOLD_ to offline cognition (N = 44; two extreme mean_BOLD_ outliers were detected and removed that did not appear in the SD_BOLD_ models above); this model revealed no robust latent effect (permuted *p* = .13). Beyond the fact that the latent permutation test did not meet threshold, we further demonstrate that the spatial representation of the mean_BOLD_ effect was almost absent at a standard spatial threshold (BSR > abs(3.00); Fig S5). Critically, no visual cortical regions were sensitive to offline cognition. Finally, to circumvent any bias in comparing thresholded SD_BOLD_ and mean_BOLD_ model results, we also correlated the raw brain weights (from the *V* singular vector; see Methods) to get a whole-brain comparison of the overlap between brain metrics. We found that each brain metric was represented by nearly orthogonal spatial representations overall (Figure 7).

**Figure 7:**
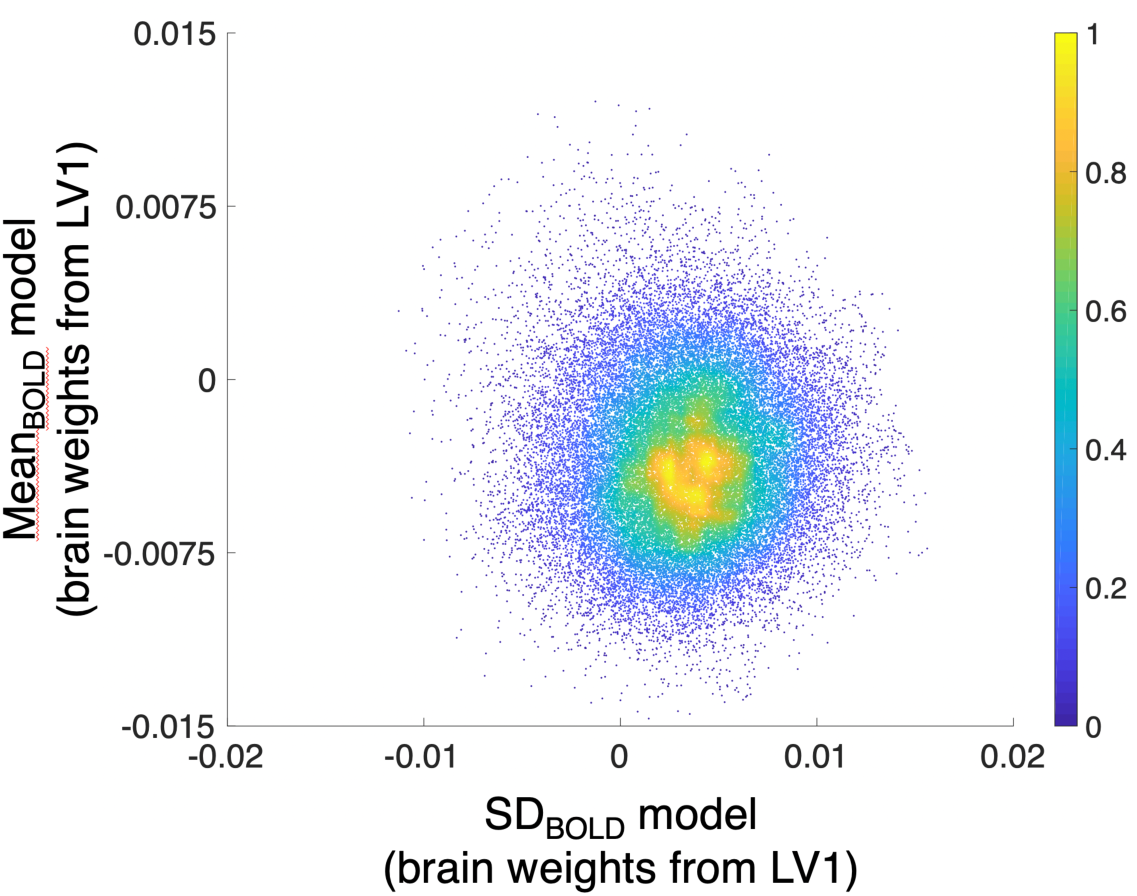
Lack of relation between SD_BOLD_ and mean_BOLD_brain weights from the first latent variable from each BOLD measure-specific PLS model.

## DISCUSSION

In the current study of healthy older adults, we found that moment-to-moment BOLD variability was an excellent differentiator of task conditions (faces and houses), and that greater upregulation of signal variability from face to house was exhibited by higher performers on a series of offline visuo-cognitive tasks. Notably, VBD was much more sensitive than mean_BOLD_-based face/house differentiation to offline behavior (even though the strict contrast of face and house conditions was largely as expected in the current data). These findings highlight variability-based differentiation (VBD) as a primary target in future work investigating the behavioral consequences of category differentiation in the (aging) human brain. Further, analyses of lower-level face and house stimulus properties (and subsequent HMAX computational model results) indicated that house images were much more differentiated in the current study. Critically, the more a subject upregulated SD_BOLD_ in line with more differentiated image content (i.e., in houses more than faces), the better their visuo-cognitive performance. Notably, this effect was most prominent in V1/V2, reflecting associated HMAX layers within which houses were detected as more feature-rich relative to faces. Combined, our results lend further support to the principled examination of temporal fluctuations in region-specific brain signals (and VBD in particular) for understanding human cognition.

### Various bases for elevated brain signal variability in response to more feature-rich visual input

At least since the 1950s, various researchers have suggested that early visual regions actively work to reduce redundancy in stimulus content (e.g., via “redundancy filtering” (Van Hateren 1992, 1993)). In accord with such theories, the visual cortex often exhibits rapid ensemble-level suppression to more common (or expected) stimuli or stimulus trains, and greater ensemble-level response to more unexpected (or complex) stimuli (Hamm and Yuste 2016; Homann *et al*. 2017; Vinken *et al*. 2017). This has important implications for BOLD variability. Because we examined within-voxel temporal variability across trials and blocks for each stimulus category, and each voxel is a moment-to-moment ensemble-level approximation of synaptic/input activity over several million neurons, lower dynamic range of BOLD in response to more predictable/homogeneous stimulus categories such as faces may indeed reflect category-level ensemble synaptic/input suppression (or in general, “coding efficiency”). Conversely, more differentiated visual input (as for houses) should ensure that dynamic range of brain responses remains broader across the stimulus period. Indeed, Hermundstad et al. (2014) argued that within-stimulus variance itself can *drive* salience, and that the visual system should thus upregulate allocated resources (perhaps expressed as dynamic range) to effectively encode more differentiated types of stimuli. The authors also showed that this effect occurs even via natural scene viewing in the absence of a specific task, similar to the use of passive viewing in the current fMRI study.

Notably, our findings also converge with computational and animal work (membrane potentials, spiking) suggesting that “perceptual uncertainty” can be probabilistically encoded by the variability of neural responses in visual cortex (Orban *et al*. 2016). In particular, Orbán et al. predict that more differentiated images should indeed yield wider distributions of neural responses. They show, for example, that the variability of V1 responses over different types of stimuli (so-called “signal variability”) increases with increasing sensory evidence (captured by contrast in the author’s study). In the present study, stimuli that were more feature-rich (i.e., houses) also yielded increased brain signal variability. Critically however, we show that individual differences in responses to such stimulus input differences are related to trait-level behavior; older adults who expressed greater upregulation of BOLD signal variability in line with sensory differentiation (i.e., houses more than faces) performed better across a battery of offline tasks.

The Orbán et al. findings provide a potential bottom-up conceptualization of this general phenomenon (i.e., signal variability as a function of perceptual input). However, top-down effects may also be important in relation to the current phenomena. In particular, predictive coding models also offer a number of potential points of insight into the current results. Conceptualizations of predictive coding (Mlynarski and Hermundstad 2018) generally presume that when neural priors for stimulus input are more “certain” (i.e., distributionally narrow), incoming stimuli within that category will also naturally be less “surprising” for the system, thus maintaining an already stable prior and making processing and prediction of future stimuli relatively easy. However, Mlynarski et al. make a critical distinction regarding uncertainty: “The degree to which incoming stimuli are surprising to the observer depends on two factors: the average surprise of the stimuli themselves, which is a property intrinsic to the stimulus distribution, and the alignment of the observer’s belief with this stimulus distribution” (p. 4). Regarding the former, the authors then go on to state that: “High-variance stimuli will therefore be more surprising to an observer, on average, than low-variance stimuli” (p. 4-5). It is this former type of uncertainty that seems most relevant to the current study. Faces are an excellent candidate for consideration as a relatively certain distribution/prior (Brodski et al. 2015; Brodski-Guerniero et al. 2017). Humans are considered face experts in general relative to other stimulus categories (Carey and Diamond 1977; Carey 1992), and faces can be reduced to a limited number of statistical dimensions, yet still be processed, discriminated, and recognized (Sirovich and Kirby 1987; O’toole *et al*. 1993; Tsao and Livingstone 2008). Accordingly, low level visual feature differentiation also appeared particularly minimized for faces in our results, potentially heightening the chance that faces remain more distributionally “certain” overall (or relatively non-updatable) for the brain, particularly in V1/V2. Indeed, Mlynarski et al. anticipate that the brain should be able to reduce neural dynamic range when processing such “reducible” types of stimuli. In our data, this idea may be reflected in a relative dampening of the dynamic range of BOLD signal for faces in higher performers.

However, house stimuli appear to be a richer and more differentiated stimulus category. Accordingly, one’s prior for houses could conceivably also be relatively broad. Humans may know generally what constitutes a “house,” but the compositional variation among house types is intuitively larger than for faces. Although predictive coding would assume that a relatively broad prior distribution for houses should adjust more rapidly to exposure to a surprising house, we would argue that it is unlikely that any natural prior distribution for houses will achieve the “certainty” humans express for faces. The brain may simply maintain broader priors for stimulus categories that are naturally more differentiated, and as predicted by Mlynarski and Hermundstad (2018), the brain may require a greater dynamic range to process such stimuli. Because greater upregulation of SD_BOLD_ for houses was indicative of better offline performance on most tasks examined, this suggests that maintaining similar neural dynamic ranges across stimulus categories (despite naturally different levels of sensory evidence) is an ineffective operational mode for the human brain. An effective brain may in fact need to continue to sample a dynamic and differentiated world (when required) to remain “optimal.” Friston et al. (2012) make a key related argument, that the brain should not fix its solutions too rigidly, instead allowing for maintenance of neural itinerancy (or instability) to approximate or maintain Bayes-optimal perception. Although fitting narrow solutions for relatively undifferentiated stimulus categories may be functional, the most adaptable brain should also be *meta-variable*, and modulate levels of dynamic range to prepare and/or respond to more differentiated levels of sensory input as required (see Fig 1). Relatedly, Marzen and Dedeo (2017) argue in their computational work that well-adapted organisms should be able to use both (1) a low-fidelity encoding regime wherever perceptual costs can be minimized, and (2) a high-fidelity mode during which perceptual costs increase with environmental complexity. In our study, such fidelity modes appear to be reflected in the variability of brain activity (thus fitting our notion of “meta-variability” to differential visual feature densities in high performers). This suggests that higher performers may do well because they can modulate “fidelity,” allowing them to encode key distinctions in their environment (see Fig 1).

### Limitations and next steps

In an effort to describe the relative feature richness within face and house stimuli using HMAX, we quantified median within-image layer features. Visual inspection of stimulus categories in Figs 4 and S1 highlights the presence of greater item-wise differentiation in houses compared to faces. Because SD_BOLD_ is computed across TRs and stimuli within-condition, our findings could represent a hybrid of variable responses at within- and between-item levels within each condition. However, notably large *t-*values (Figure 5) suggest that condition differences in median feature richness remain key despite item variance within conditions. Faster imaging methods (EEG/MEG) are needed in future work to disentangle the relative influence of within- and between-item differences on brain signal variability levels, despite the loss of spatial information such techniques would impose. Further, future studies could re-test the current hypotheses by using visual stimuli beyond faces and houses. Based on convergence between initial HMAX model layers and performance-relevant signal variability in V1/V2, we anticipate that the current effects would likely hold using any image sets that differ reliably in feature differentiation in V1/V2. An alternative and strong test of the current findings would involve constructing image sets with maximal differences in feature density that are detectable in layers beyond V1/V2; this would then allow an examination of whether brain signal variability is also maximally differentiated at later stages of the ventral visual stream given such stimuli, particularly in higher performing adults. Finally, the current sample contains only older adults; although this serves as an effective control against baseline age differences in SD_BOLD_ driving any associated behavioral effects (see Garrett et al., 2013), future work could re-test the current hypotheses within different age groups (e.g., young adults only) to verify the generality of the present effects.

### Conclusion

We conclude that the ability to align brain signal variability (especially in primary visual cortex) to the complexity of visual input may mark heightened trait-level behavioral performance in older adults.

## Acknowledgements

D.D.G was supported by an Emmy Noether Programme grant from the German Research Foundation. U.L. acknowledges financial support from the Intramural Innovation Fund of the Max Planck Society. D.D.G and U.L. were also partially supported by the Max Planck UCL Centre for Computational Psychiatry and Ageing Research. Finally, we would like to thank Martin Hebart for helpful discussions regarding implementation of HMAX.

## SUPPLEMENTARY MATERIAL

### Results

#### HMAX results

**Figure S1:**
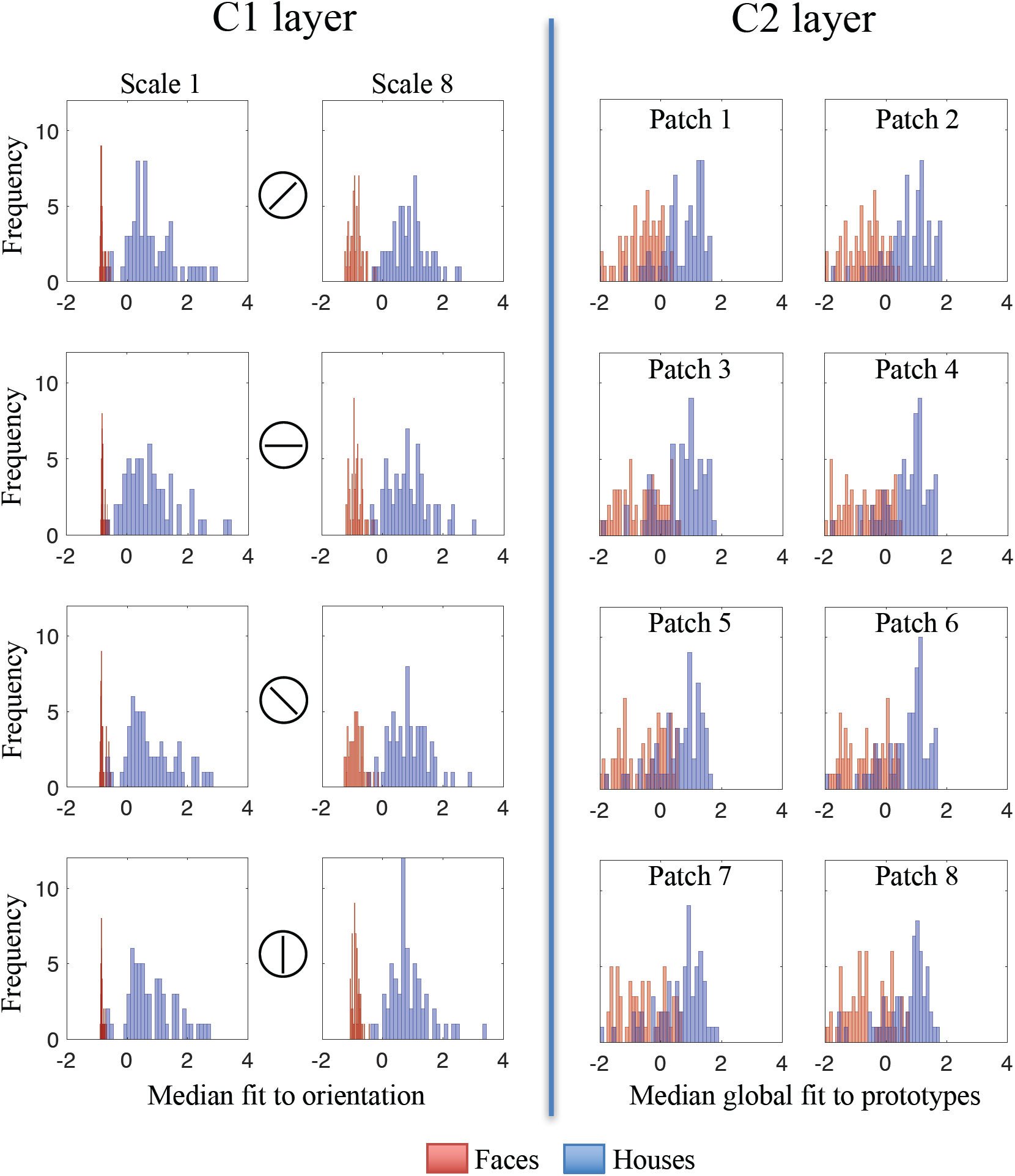
HMAX C1 (simple feature) and C2 (composite feature) distributions for face and house stimuli. All 60 unique images per condition from the fMRI stimulus set were submitted to the model. At the C1 layer (left), the median of each C1 map is shown, averaging over all locations within-image, for the smallest and largest receptive field sizes (i.e., scale bands 1 and 8) and each spatial orientation separately. At the C2 layer (right), the median within-image fit to a library of 400 feature prototypes is displayed for each of eight neighborhood (receptive field. Or “patch”) sizes. Higher values for C1 and C2 indicate higher “feature richness.” All C1 and C2 value ranges (x-axes) are z-normalized.

##### SDBOLD model results

###### Group level multivariate model of face-house modulation of SD_BOLD_

We examined whether a group level rise in brain signal variability was present on the house relative to the face condition. Indeed, we noted a significant latent variable (permuted *p* = .005) expressing that house stimuli generated more variable brain signal responses than face stimuli (see Figure S2). Conversely, there was one small cluster (in blue) in lateral occipital cortex in which temporal variability was higher during the face condition.

**Figure S2:**
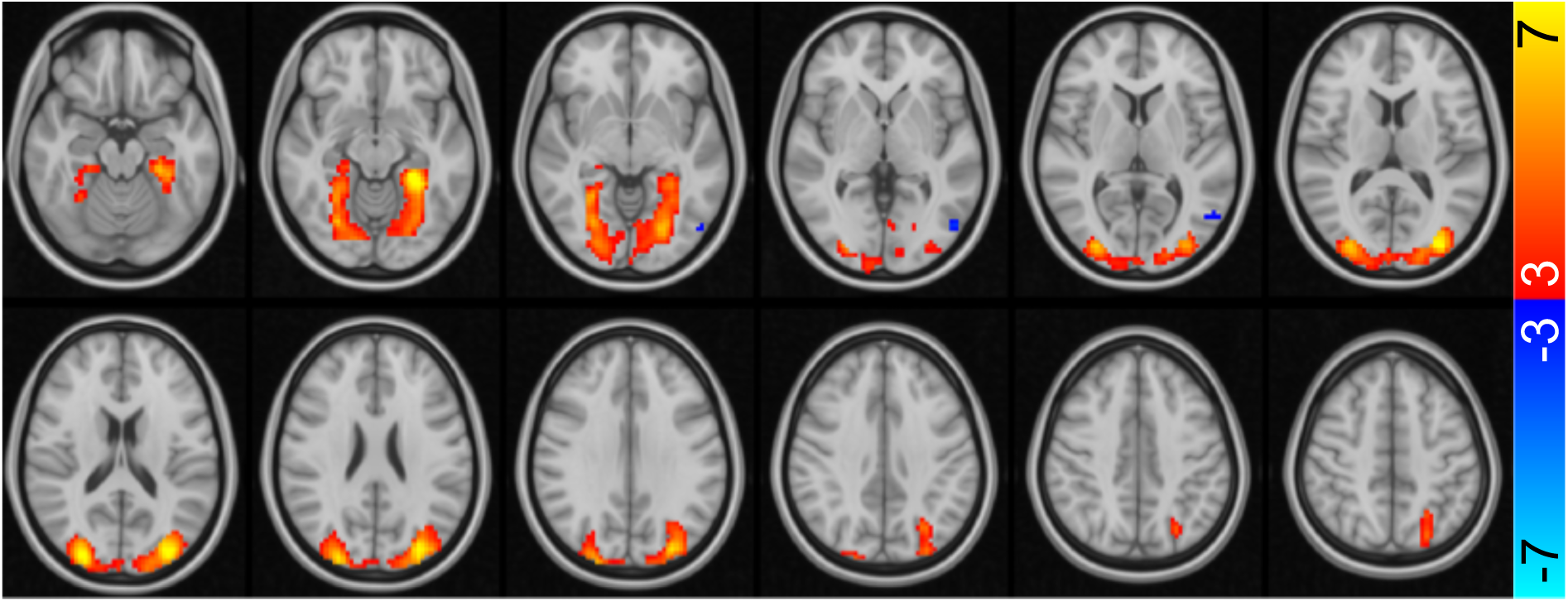
Task PLS model differentiating SD_BOLD_ in face and house conditions. SD_BOLD_ was typically higher in the house condition (red regions) compared to the face condition (single blue cluster). Color bar = bootstrap ratio ranges. Slices increase from Z=-18 (top left) in 6 mm increments.

###### Behavioural PLS model linking SD_BOLD_ modulations to offline cognition

In the main results, we showed that SD_BOLD_ upregulation from face to house was greater in high performers at the latent level (Figure 6). Because this effect collapses across cognitive measures within the model, here we show the correlation value between each cognitive measure and the overall brain score depicted in Figure 6. This latent effect was bootstrap robust (bootstrap CIs do not cross zero) for 3/4 (accuracy), 8/8 (RT_mean_), and 7/8 (RT_SD_) cognitive performance measures (see Figure S3).

**Figure S3:**
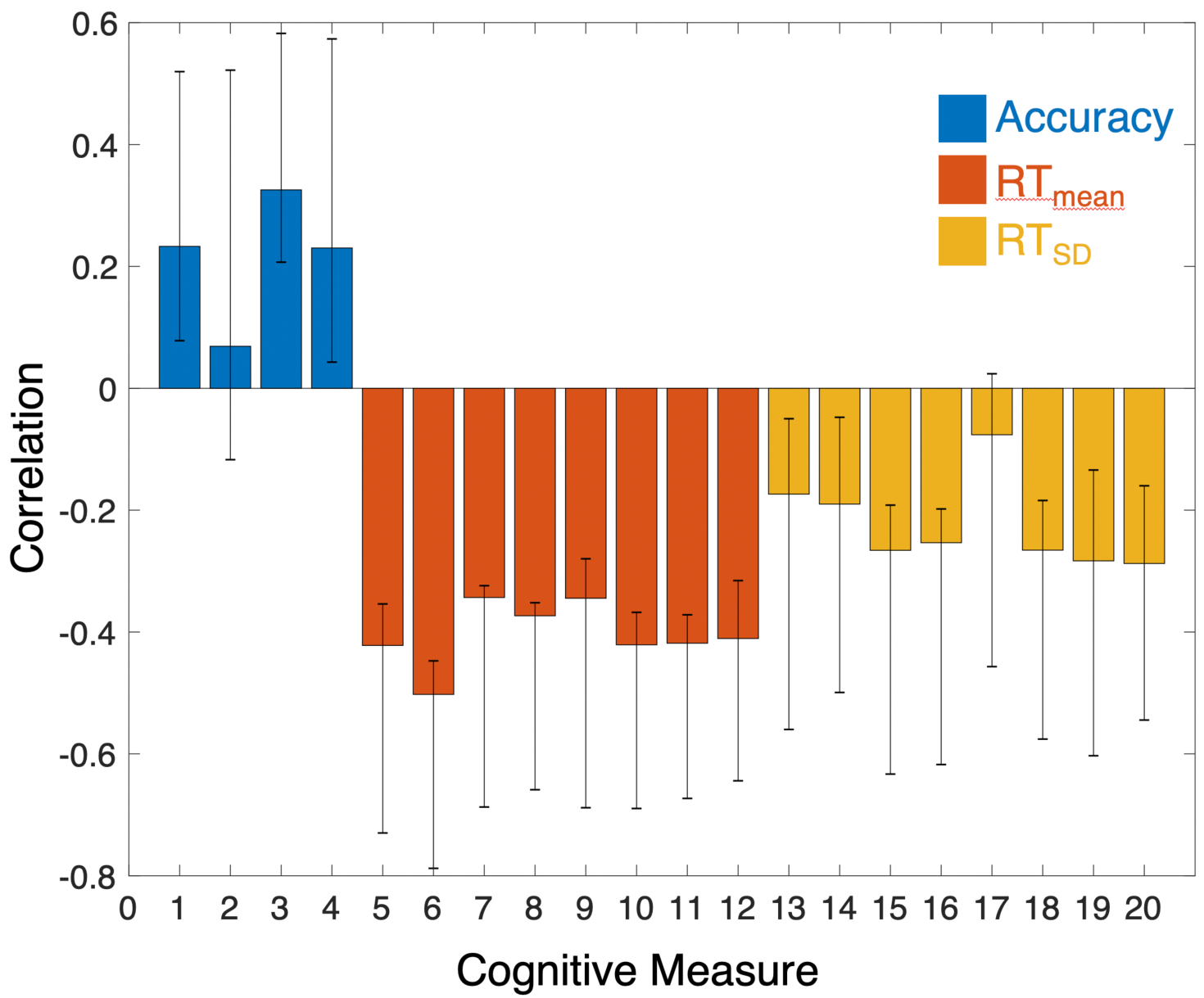
Correlation between individual cognitive measures and the overall brain score depicted in Figure 6 (main results). The measures are numbered as follows: (1) Letter updating; (2) Spatial updating; (3) 2-back; (4) Raven’s progressive matrices; (5 & 6) Stroop congruent and incongruent; (7 & 8) Number/letter switch and non-switch; (9 & 10) Global/local switch and non-switch; (11 & 12) Face/word switch and nonswitch. Measures 13-20 represent the same tasks as for 5-12, but RT_SD_ instead of RT_mean_.

**Table S1:**
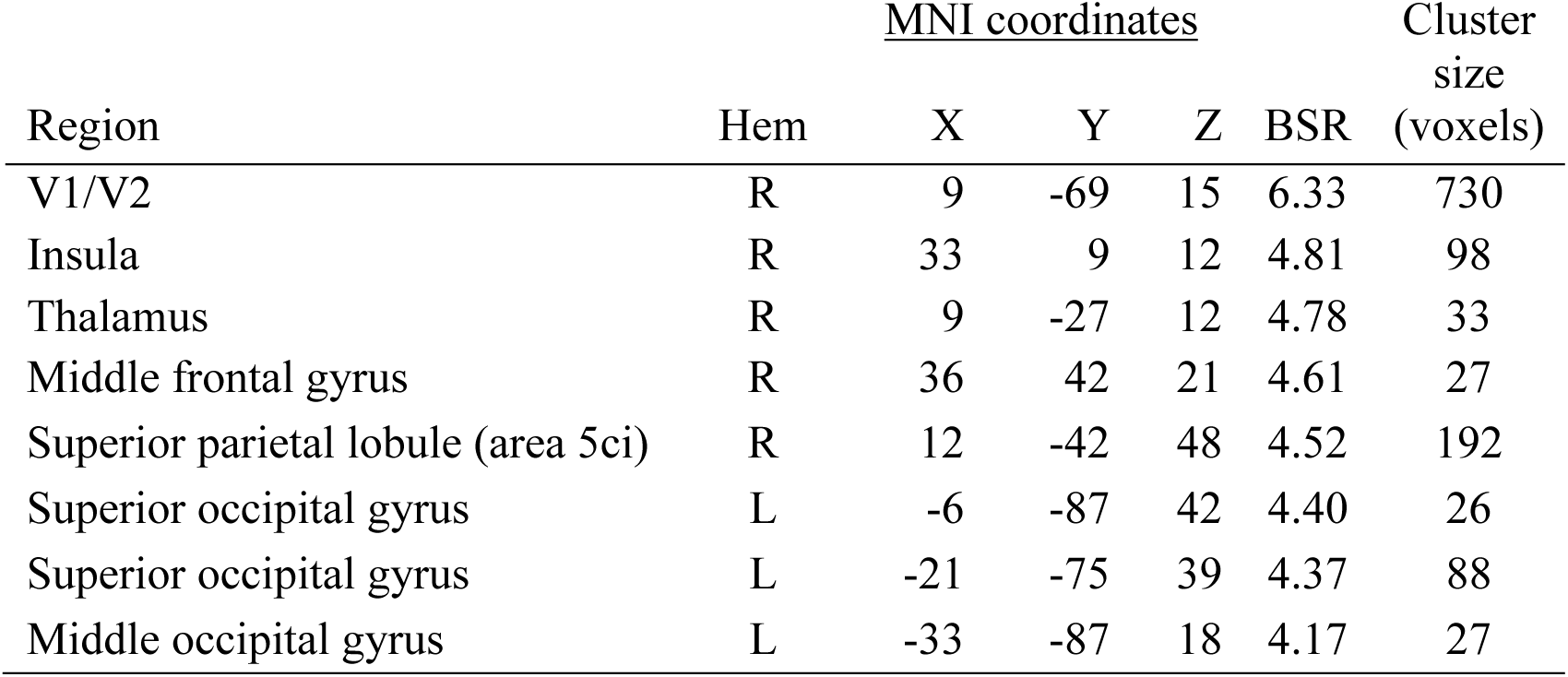
Multivariate PLS model peak activations, bootstrap ratios, and cluster sizes: Regions expressing heightened SD_BOLD_ on house vs face in relation to faster and more stable offline speeded performance. SD = standard deviation; BOLD = blood oxygen level-dependent; Hem = hemisphere; MNI = Montreal Neurological Institute; BSR = bootstrap ratio (model salience/bootstrapped standard error).

| Region | Hem | <u>MNI coordinates</u> |  |  | BSR | Cluster size (voxels) |
| --- | --- | --- | --- | --- | --- | --- |
|  |  | X | Y | Z |  |  |
| V1/V2 | R | 9 | -69 | 15 | 6.33 | 730 |
| Insula | R | 33 | 9 | 12 | 4.81 | 98 |
| Thalamus | R | 9 | -27 | 12 | 4.78 | 33 |
| Middle frontal gyrus | R | 36 | 42 | 21 | 4.61 | 27 |
| Superior parietal lobule (area 5ci) | R | 12 | -42 | 48 | 4.52 | 192 |
| Superior occipital gyrus | L | -6 | -87 | 42 | 4.40 | 26 |
| Superior occipital gyrus | L | -21 | -75 | 39 | 4.37 | 88 |
| Middle occipital gyrus | L | -33 | -87 | 18 | 4.17 | 27 |

##### Mean_BOLD_ model results

**Figure S4:**
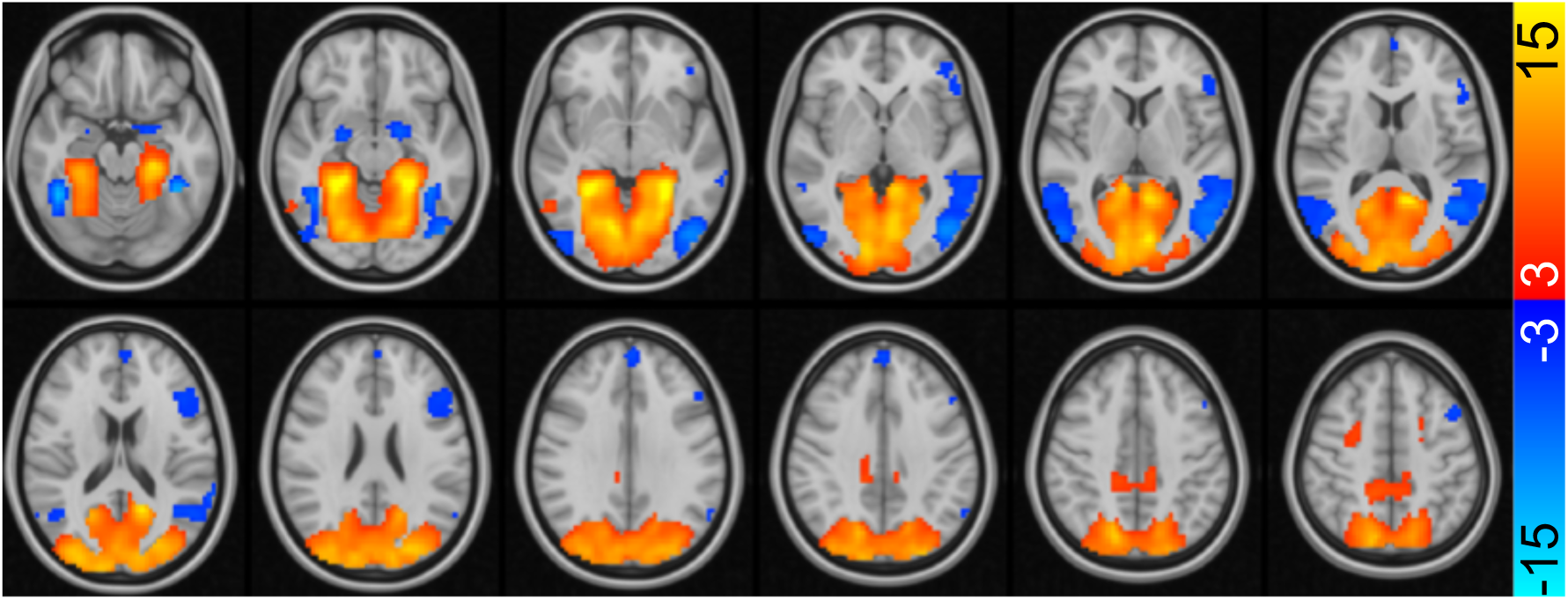
Mean_BOLD_ contrast between faces (higher in blue regions) and houses (higher in red regions). Slices increase from Z=-18 (top left) in 6 mm increments.

**Figure S5:**
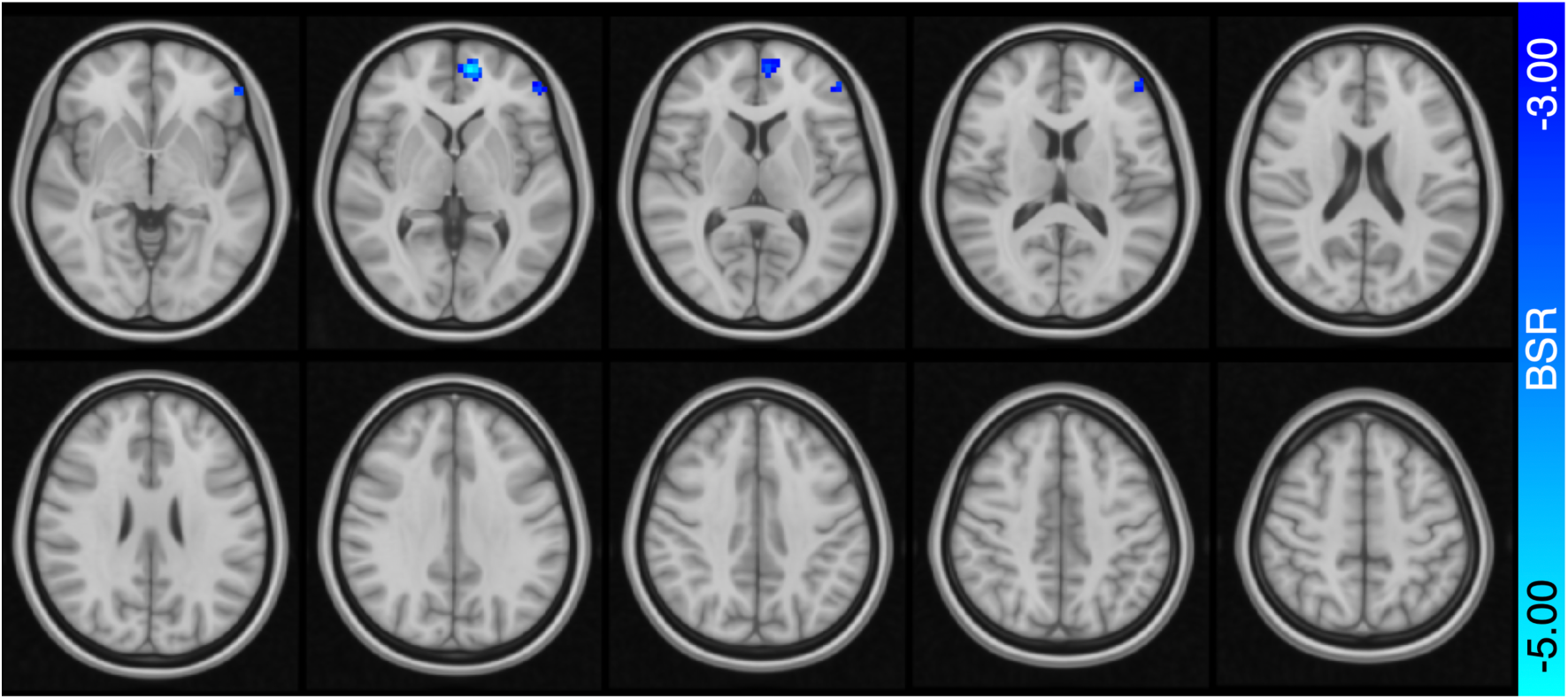
Spatial pattern from behavioural PLS model linking mean_BOLD_ to offline cognitive performance. No significant latent variable was found linking brain and behavior (permuted *p* = .13); however, for completeness, we show that modest PFC clusters denoted higher mean_BOLD_ in the face vs. house condition in higher offline performers (latent *r =* .49). Slices increase from Z=-4 (top left) in 6 mm increments.

## REFERENCES

Beck JM, Ma WJ, Kiani R, Hanks T, Churchland AK, Roitman J, Shadlen MN, Latham PE, Pouget A. 2008. Probabilistic population codes for Bayesian decision making. Neuron 60:1142–1152.

Beckmann CF, Smith SM. 2004. Probabilistic independent component analysis for functional magnetic resonance imaging. IEEE Trans Med Imaging 23:137–152.

Brodski A, Paasch GF, Helbling S, Wibral M. 2015. The Faces of Predictive Coding. J Neurosci 35:8997–9006.

Brodski-Guerniero A, Paasch GF, Wollstadt P, Ozdemir I, Lizier JT, Wibral M. 2017. Information-Theoretic Evidence for Predictive Coding in the Face-Processing System. J Neurosci 37:8273–8283.

Carey S. 1992. Becoming a face expert. Philos Trans R Soc Lond B Biol Sci 335:95–102; discussion 102-103.

Carey S, Diamond R. 1977. From piecemeal to configurational representation of faces. Science 195:312–314.

Carp J, Gmeindl L, Reuter-Lorenz PA. 2010. Age differences in the neural representation of working memory revealed by multi-voxel pattern analysis. Front Hum Neurosci 4:217.

Efron B, Tibshirani RJ. 1993. An Introduction to the Bootstrap. lBoston, MA: Springer US.

Friston K, Breakspear M, Deco G. 2012. Perception and self-organized instability. Front Comput Neurosci 6:44.

Garrett D, Epp S, Perry A, Lindenberger U. 2018. Local temporal variability reflects functional network integration in the human brain: On the crucial role of the thalamus. BioRxiv:184739.

Garrett DD, Kovacevic N, McIntosh AR, Grady CL. 2010. Blood oxygen level-dependent signal variability is more than just noise. J Neurosci 30:4914–4921.

Garrett DD, Kovacevic N, McIntosh AR, Grady CL. 2011. The importance of being variable. J Neurosci 31:4496–4503.

Garrett DD, Kovacevic N, McIntosh AR, Grady CL. 2013. The modulation of BOLD variability between cognitive states varies by age and processing speed. Cereb Cortex 23:684–693.

Garrett DD, Lindenberger U, Hoge RD, Gauthier CJ. 2017. Age differences in brain signal variability are robust to multiple vascular controls. Sci Rep 7:10149.

Garrett DD, McIntosh AR, Grady CL. 2014. Brain signal variability is parametrically modifiable. Cereb Cortex 24:2931–2940.

Garrett DD, Nagel IE, Preuschhof C, Burzynska AZ, Marchner J, Wiegert S, Jungehulsing GJ, Nyberg L, Villringer A, Li SC, Heekeren HR, Backman L, Lindenberger U. 2015. Amphetamine modulates brain signal variability and working memory in younger and older adults. Proc Natl Acad Sci U S A 112:7593–7598.

Garrett DD, Samanez-Larkin GR, MacDonald SW, Lindenberger U, McIntosh AR, Grady CL. 2013. Moment-to-moment brain signal variability: a next frontier in human brain mapping? Neurosci Biobehav Rev 37:610–624.

Grady CL, Garrett DD. 2014. Understanding variability in the BOLD signal and why it matters for aging. Brain Imaging Behav 8:274–283.

Hamm JP, Yuste R. 2016. Somatostatin Interneurons Control a Key Component of Mismatch Negativity in Mouse Visual Cortex. Cell Rep 16:597–604.

Hermundstad AM, Briguglio JJ, Conte MM, Victor JD, Balasubramanian V, Tkacik G. 2014. Variance predicts salience in central sensory processing. Elife 3:308.

Homann J, Koay SA, Glidden AM, Tank DW, Berry MJ. 2017. Predictive Coding of Novel versus Familiar Stimuli in the Primary Visual Cortex. BioRxiv.

Jenkinson M, Beckmann CF, Behrens TE, Woolrich MW, Smith SM. 2012. Fsl. Neuroimage 62:782–790.

Kinchla RA, Solis-Macias V, Hoffman J. 1983. Attending to different levels of structure in a visual image. Percept Psychophys 33:1–10.

Kleemeyer MM, Kuhn S, Prindle J, Bodammer NC, Brechtel L, Garthe A, Kempermann G, Schaefer S, Lindenberger U. 2016. Changes in fitness are associated with changes in hippocampal microstructure and hippocampal volume among older adults. Neuroimage 131:155–161.

Kleemeyer MM, Polk TA, Schaefer S, Bodammer NC, Brechtel L, Lindenberger U. 2017. Exercise-Induced Fitness Changes Correlate with Changes in Neural Specificity in Older Adults. Front Hum Neurosci 11:123.

Knill DC, Pouget A. 2004. The Bayesian brain: the role of uncertainty in neural coding and computation. Trends Neurosci 27:712–719.

Krishnan A, Williams LJ, McIntosh AR, Abdi H. 2011. Partial Least Squares (PLS) methods for neuroimaging: a tutorial and review. Neuroimage 56:455–475.

Ma WJ, Beck JM, Latham PE, Pouget A. 2006. Bayesian inference with probabilistic population codes. Nat Neurosci 9:1432–1438.

Marzen SE, DeDeo S. 2017. The evolution of lossy compression. J R Soc Interface 14:20170166.

McIntosh AR, Bookstein FL, Haxby JV, Grady CL. 1996. Spatial pattern analysis of functional brain images using partial least squares. Neuroimage 3:143–157.

McKeeff TJ, Tong F. 2007. The timing of perceptual decisions for ambiguous face stimuli in the human ventral visual cortex. Cereb Cortex 17:669–678.

Minear M, Park D. 2004. A lifespan database of adult facial stimuli. Behavior Research Methods, Instruments, & Computers:630–633.

Mlynarski WF, Hermundstad AM. 2018. Adaptive coding for dynamic sensory inference. Elife 7.

Mooney CM, Ferguson GA. 1951. A new closure test. Can J Psychol 5:129–133.

O’Toole AJ, Deffenbacher KA, Valentin D, Abdi H. 1993. Low-dimensional representation of faces in higher dimensions of the face space. Journal of the Optical Society of America A 10:405.

Orban G, Berkes P, Fiser J, Lengyel M. 2016. Neural Variability and Sampling-Based Probabilistic Representations in the Visual Cortex. Neuron 92:530–543.

Park J, Carp J, Hebrank A, Park DC, Polk TA. 2010. Neural specificity predicts fluid processing ability in older adults. J Neurosci 30:9253–9259.

Raven J, Raven JC, Court JH. 1998. Manual for Raven’s progressive matrices and vocabulary scales. Section 1:General overview. Oxford, UK: Oxford Psychologists Press.

Riesenhuber M, Poggio T. 1999. Hierarchical models of object recognition in cortex. Nat Neurosci 2:1019–1025.

Schmiedek F, Hildebrandt A, Lovden M, Lindenberger U, Wilhelm O. 2009. Complex span versus updating tasks of working memory: the gap is not that deep. J Exp Psychol Learn Mem Cogn 35:1089–1096.

Schmiedek F, Lovden M, Lindenberger U. 2010. Hundred Days of Cognitive Training Enhance Broad Cognitive Abilities in Adulthood: Findings from the COGITO Study. Front Aging Neurosci 2.

Schmitter-Edgecombe M, Langill M. 2006. Costs of a predictable switch between simple cognitive tasks following severe closed-head injury. Neuropsychology 20:675–684.

Serre T, Kouh M, Cadieu C, Knoblich U, Kreiman G, Poggio T. 2005. A Theory of Object Recognition: Computations and Circuits in the Feedforward Path of the Ventral Stream in Primate Visual Cortex. AI Memo 2005-036/CBCL Memo 259

Serre T, Oliva A, Poggio T. 2007. A feedforward architecture accounts for rapid categorization. Proc Natl Acad Sci U S A 104:6424–6429.

Serre T, Wolf L, Bileschi S, Riesenhuber M, Poggio T. 2007. Robust object recognition with cortex-like mechanisms. IEEE Trans Pattern Anal Mach Intell 29:411–426.

Sirovich L, Kirby M. 1987. Low-dimensional procedure for the characterization of human faces. J Opt Soc Am A 4:519–524.

Smith AM, Lewis BK, Ruttimann UE, Ye FQ, Sinnwell TM, Yang Y, Duyn JH, Frank JA. 1999. Investigation of low frequency drift in fMRI signal. Neuroimage 9:526–533.

Smith SM, Jenkinson M, Woolrich MW, Beckmann CF, Behrens TE, Johansen-Berg H, Bannister PR, De Luca M, Drobnjak I, Flitney DE, Niazy RK, Saunders J, Vickers J, Zhang Y, De Stefano N, Brady JM, Matthews PM. 2004. Advances in functional and structural MR image analysis and implementation as FSL. Neuroimage 23 Suppl 1:S208–219.

Stroop JR. 1935. Studies of interference in serial verbal reactions. Journal of Experimental Psychology 18:643–662.

Tsao DY, Livingstone MS. 2008. Mechanisms of face perception. Annu Rev Neurosci 31:411–437.

van Hateren JH. 1992. A theory of maximizing sensory information. Biol Cybern 68:23–29.

Van Hateren JH. 1993. Spatiotemporal contrast sensitivity of early vision. Vision Res 33:257–267.

Vinken K, Vogels R, Op de Beeck H. 2017. Recent Visual Experience Shapes Visual Processing in Rats through Stimulus-Specific Adaptation and Response Enhancement. Curr Biol 27:914–919.

Wen H, Shi J, Zhang Y, Lu KH, Cao J, Liu Z. 2017. Neural Encoding and Decoding with Deep Learning for Dynamic Natural Vision. Cereb Cortex:1–25.

Yeung N, Nystrom LE, Aronson JA, Cohen JD. 2006. Between-task competition and cognitive control in task switching. J Neurosci 26:1429–1438.

